# Evaluation of sequencing strategies for whole-genome imputation with hybrid peeling

**DOI:** 10.1101/824631

**Authors:** Roger Ros-Freixedes, Andrew Whalen, Gregor Gorjanc, Alan J Mileham, John M Hickey

**Author notes:** Email addresses: AW, GG, AJM, JMH.

## Abstract

**Background:** For assembling large whole-genome sequence datasets to be used routinely in research and breeding, the sequencing strategy should be adapted to the methods that will later be used for variant discovery and imputation. In this study we used simulation to explore the impact that the sequencing strategy and level of sequencing investment have on the overall accuracy of imputation using hybrid peeling, a pedigree-based imputation method well-suited for large livestock populations.

**Methods:** We simulated marker array and whole-genome sequence data for fifteen populations with simulated or real pedigrees that had different structures. In these populations we evaluated the effect on imputation accuracy of seven methods for selecting which individuals to sequence, the generation of the pedigree to which the sequenced individuals belonged, the use of variable or uniform coverage, and the trade-off between the number of sequenced individuals and their sequencing coverage. For each population we considered four levels of investment in sequencing that were proportional to the size of the population.

**Results:** Imputation accuracy largely depended on pedigree depth. The distribution of the sequenced individuals across the generations of the pedigree underlay the performance of the different methods used to select individuals to sequence. Additionally, it was critical to balance high imputation accuracy in early generations as well as in late generations. Imputation accuracy was highest with a uniform coverage across the sequenced individuals of around 2x rather than variable coverage. An investment equivalent to the cost of sequencing 2% of the population at 2x provided high imputation accuracy. The gain in imputation accuracy from additional investment diminished with larger populations and larger levels of investment. However, to achieve the same imputation accuracy, a proportionally greater investment must be used in the smaller populations compared to the larger ones.

**Conclusions:** Suitable sequencing strategies for subsequent imputation with hybrid peeling involve sequencing around 2% of the population at a uniform coverage around 2x, distributed preferably from the third generation of the pedigree onwards. Such sequencing strategies are beneficial for generating whole-genome sequence data in populations with deep pedigrees of closely related individuals.

## Background

The coupling of appropriate sequencing strategies and powerful imputation methods enables the generation of large datasets of sequenced individuals at a low cost. In this paper we assess the impact that the sequencing strategy and the level of investment in sequencing have on the imputation of whole-genome sequence data for entire livestock populations with a pedigree-based imputation method that does not require haplotype reference panels, such as hybrid peeling [1].

Sequence data has the potential to empower the identification of causal variants that underlie quantitative traits or diseases [2–5], enhance livestock breeding [6–8], and increase the precision and scope of population genetic studies [9,10]. For sequence data to be used routinely in research and breeding, low-cost sequencing strategies must be deployed in order to assemble large data sets that capture most of the genetic diversity in a population and enable harnessing of its potential. A range of low-cost sequencing strategies have been proposed, which involve sequencing a subset of individuals and then performing imputation of whole-genome sequence data for the remaining individuals. Possible sequencing strategies range from sequencing key ancestors in a population at high coverage [3,7] to sequencing a large subset of the individuals in a population at low coverage [11–13].

For the implementation of these sequencing strategies, several methods have been proposed to select the individuals to sequence and the coverage at which to sequence them in both human and livestock populations. Some methods are based on pedigree information only, while others are based on genomic information. Methods based on pedigree information include the method developed by Boichard [14], which was originally intended for analysing pedigrees of large populations and iteratively selects the ancestors that contribute the greatest pedigree-inferred marginal contributions, as well as methods proposed by Druet et al. [7], which maximise either the genetic relationship between the sequenced individuals and the rest of the population or, on the contrary, the number of independent genomes captured. The approaches of the methods based on genomic information are diverse, but these methods typically use haplotype libraries derived from phased marker array genotypes [7,15–19] or the genomic relationship matrix [20,21].

Some of these methods have been designed to select sets of individuals for producing haplotype reference panels. Many imputation methods, like the population-based imputation algorithms traditionally used in human genetics, use these reference panels to find haplotypes that match those of the imputed individuals [22–27]. In livestock, these reference panels are typically developed by sequencing key ancestors, which are expected to carry many of the high-frequency haplotypes in the population, at high coverage. However, alternative imputation methods exist that do not use reference panels and it is unclear whether the methods that have been proposed to select the individuals to sequence are well-suited for them.

We have recently proposed the use of ‘hybrid peeling’ [1], a fast and accurate imputation method explicitly designed for jointly calling, phasing and imputing whole-genome sequence data in large and complex multi-generational pedigreed populations where individuals can be sequenced at variable coverage or not sequenced at all. Hybrid peeling is a two-step process. In the first step, multi-locus iterative peeling is performed to estimate the segregation probabilities for a subset of segregating sites (e.g., the markers on a genotyping array). In the second step, the segregation probabilities are used to perform fast single-locus iterative peeling on every segregating site discovered in the genome. This two-step process allows the computationally demanding multi-locus peeling step to be performed on only a subset of the variants, while still leveraging linkage information for the remaining variants. We have recently shown that hybrid peeling is a powerful method for imputing whole-genome sequence data from marker array data in large pedigreed livestock populations where only a small fraction of individuals need to be sequenced, mostly at low coverage [28]. We have also evaluated the effect of the factors that determine individual-wise and variant-wise imputation accuracy [28].

Our hypothesis is that the choice of sequencing strategy and method for selecting which individuals to sequence should be adapted to the methods that will later be used for variant discovery and for imputation. A body of work already exists for the design of reference panels for population-based imputation methods [7,19,25,29–31]. To our knowledge, a similar body of work is absent for pedigree-based imputation methods that do not require haplotype reference panels, such as hybrid peeling.

The objectives of this study were to assess: (i) the performance of hybrid peeling based on the structure of the pedigree and the level of investment in sequencing; and (ii) the performance of hybrid peeling based on the choice of sequencing strategy. For this purpose we simulated whole-genome sequence data under different sequencing strategies for a series of simulated and real pedigrees. We assessed the effect of six factors on the population-wide imputation accuracy of hybrid peeling: (i) pedigree structure and population size; (ii) level of investment in sequencing; (iii) the method for selecting individuals to sequence; (iv) the generation of the pedigree to which the sequenced individuals belonged; (v) the use of variable or uniform sequencing coverage; and (vi) the trade-off between the number of sequenced individuals and the sequencing coverage. Our results show that hybrid peeling generally performs well regardless of the sequencing strategy as long as the number of sequenced individuals is maximised for the available budget and distributed widely across the generations of the pedigree. We found that sequencing individuals at uniform coverage was at least as powerful as sequencing individuals at variable coverage. We conclude with recommendations on sequencing strategies for population-wide studies with limited budgets.

## Materials and Methods

We simulated marker array genotype data and whole-genome sequence data for twelve populations with simulated pedigrees and three populations with real pedigrees. For each population we defined four different levels of investment in sequencing, proportional to population size and equivalent to sequencing between 0.5% and 5% of the population at 2x. We structured the analysis of the data into four tests. In Test 1 we used different methods for selecting the individuals to sequence and the coverage at which they were sequenced in order to assess the suitability of these methods when imputation was performed with hybrid peeling. In Test 2 we focused all sequencing into specific generations of the pedigree and assessed the effect that the generation to which the sequenced individuals belonged had on imputation accuracy. In Test 3 we compared the imputation accuracy achieved by sequencing the same set of individuals at either variable or uniform coverage. In Test 4 we sequenced different numbers of individuals at different levels of coverage and quantified the trade-off between the number of sequenced individuals and the coverage at which they are sequenced. In what follows we describe in detail how the data was generated and how the different tests were performed.

### Simulated data

Using AlphaSim [32] we simulated genotype data for a total of twelve populations with simulated pedigrees and three populations with real pedigrees to represent different pedigree structures found in livestock populations. For each population we simulated marker array genotype data at two densities (high and low) based on genotyping schemes currently used in typical livestock breeding programs and we simulated whole-genome sequence data based on each sequencing strategy tested. For each scenario we simulated two replicates and the results were averaged across replicates.

#### Populations with simulated pedigrees

Pedigrees were simulated to represent populations with different numbers of generations and different generation sizes. The simulated pedigrees consisted of 2, 5, 10, or 15 discrete generations with 500, 1,000, or 2,000 individuals per generation. Thus, the size of the simulated pedigrees ranged from 1,000 to 30,000 individuals. In each generation, truncation selection on true breeding values was performed for a polygenic trait using selection proportions of 5% for sires and 25% for dams, with each selected parent contributing the same amount of progeny. The polygenic trait was simulated to have a heritability of 0.3 and was controlled by 20,000 variants, which were selected randomly from the genotype data simulated as described below and whose effects were sampled from a normal distribution. The sires and dams that produced the first generation in each scenario were also added to the pedigree and considered to be the ‘base generation’. Individuals who did not contribute progeny were not pruned. The pedigrees were sorted so that parents appeared before their progeny. A summary of the structure of the simulated pedigrees is provided in Table 1.

**Table 1.**
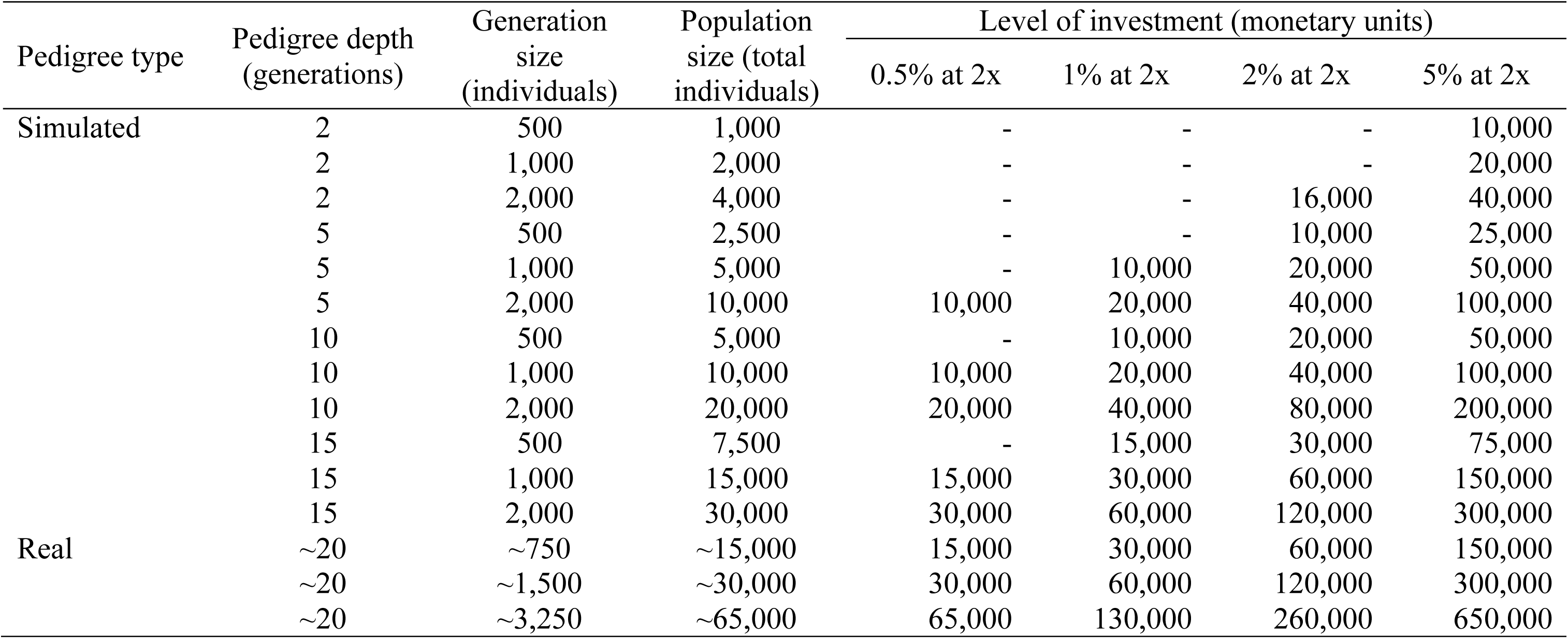
Summary of the structure of the simulated and real pedigrees and levels of investment.

Genotype data was simulated for 20 chromosomes, each 100 cM in length. For each chromosome we generated 1,000 base haplotypes assuming a chromosome length of 10^8^ base pairs, a per-site mutation rate of 2.5×10^−8^, a between-site recombination rate of 1×10^−8^, and an effective population size (N_e_) that varied over time in accordance with estimates for a livestock population [33] using the Markovian Coalescent Simulator (MaCS) [34]. We then used AlphaSim [32] to drop the base haplotypes through the simulated multi-generational pedigrees. A total of 150,000 SNPs per chromosome (3 million SNPs genome-wide) were simulated in order to represent the whole-genome sequence. A subset of 3,000 SNPs per chromosome (60,000 SNPs genome-wide) was used as a high-density marker array (HD). A smaller subset of 300 SNPs per chromosome (6,000 SNPs genome-wide) nested within the high-density marker array was used as a low-density marker array (LD). Each individual was assigned HD or LD marker array data according to the following proportions: in the base generation (parents of generation 1), 75% individuals were genotyped, of which 67% and 33% were genotyped at high and low density, respectively; in the remaining generations, 95% of individuals were genotyped, of which 15% and 85% were genotyped at high and low density, respectively. The genotyped/non-genotyped status was assigned randomly but the marker array density was assigned prioritising HD for individuals that produced the most (grand)progeny, except in the final generation, where it was assigned randomly.

#### Populations with real pedigrees

In addition to the simulated pedigrees, three real livestock pedigrees from commercial pig breeding lines (Genus PIC, Hendersonville, TN) were used. The three real pedigrees were selected to represent different population sizes: 15,187 (15k), 29,974 (30k), and 64,598 (65k) individuals. The pedigrees of each population were sorted so that parents appeared earlier than their progeny. Genotype data was simulated following the same steps used for the simulated pedigrees, but the marker array density at which each individual was genotyped was based on the density at which they were genotyped in reality. A summary of the structure of the real pedigrees is provided in Table 1.

### Levels of investment and sequencing costs

For each of the populations we tested four levels of investment in sequencing, proportional to the population size. The levels of investment corresponded to the equivalent cost of sequencing 0.5%, 1%, 2%, or 5% of the population at 2x (sequencing an individual at 2x was assumed to cost 200 monetary units, MU). Scenarios where total investment was less than 10,000 MU were ignored. The levels of investment are summarised in Table 1.

We assumed a library preparation cost of 40 MU and a linear sequencing cost between 1x and 5x of 80 MU per x. Thus, the combination of library preparation and sequencing at 1x, 2x, and 5x had respectively a cost of 120 MU, 200 MU, and 440 MU. We assumed that the combined cost scaled to 500 MU at 15x and 850 MU at 30x sequencing. These non-linear cost assumptions reflect non-linear costs that are prevalent in the current market.

### Allocation of sequencing resources

The individuals to be sequenced and their sequencing coverage were selected according to the design of each of the four tests. The number of sequenced individuals in each test scenario was determined by their sequencing coverage and the level of investment.

#### Test 1: Method for selecting individuals to sequence and coverage

We tested the effect of different methods for selecting individuals to sequence and their sequencing coverage on the imputation accuracy when using hybrid peeling. We considered the following four methods:

##### Top sires and dams (pedigree-based)

We ranked the sires and dams based on their number of genotyped progeny and grandprogeny. We split sequencing resources equally between sequencing sires (50% of the investment) and dams (the other 50%) with more progeny, referred to as ‘top sires’ and ‘top dams’. Because in livestock dams contribute fewer progeny than sires, the top sires were sequenced at 2x and the top dams at 1x. We refer to this method as ‘TopSiresAndDams’.

##### Key ancestors (pedigree-based)

We identified key ancestors of a population using PEDIG [14]. PEDIG iteratively selects the set of individuals that contribute the greatest pedigree-inferred marginal contributions. The selected individuals were sequenced at 15x to represent the sequencing strategy of key ancestors at high coverage. We refer to this method as ‘KeyAncestors’.

##### Pedigree connectedness (pedigree-based)

We selected individuals sequentially following the ‘REL’ method proposed by Druet et al. [7]. This method maximises the genetic relationship between the sequenced individuals and the rest of the population while accounting for the relationships among the sequenced individuals. We used a stepwise strategy to select, first, the individual with the highest average pedigree relationship with the rest of the population, and then, the individuals with the highest score calculated as (**A**_*s*_^-1^ **a**_*s*_)^t^ **1**_*s*_, where **A**_*s*_ is the pedigree relationship matrix between the *s* sequenced individuals and **a**_*s*_ is the vector of average pedigree relationships between each sequenced individual *s* and the rest of individuals in the population. The selected individuals were sequenced at 15x to represent a sequencing strategy of key individuals at high coverage. We refer to this method as ‘PedConnect’.

##### AlphaSeqOpt (haplotype-based)

We used the joint AlphaSeqOpt method, which has two stages. In the first stage, AlphaSeqOpt part 1 [17] was used to identify the individuals whose haplotypes represented a large proportion of the population haplotypes (referred to as ‘focal individuals’) and then variable levels of sequence coverage (i.e., not sequenced or sequenced at 1x, 2x, 5x, 15x, or 30x) were assigned to these focal individuals and their parents and grandparents so that the expected phasing accuracy of the haplotypes that they carry was maximised. In the second stage, AlphaSeqOpt part 2 [18] was used to identify individuals that carried haplotypes whose cumulative coverage was low at the end of the first stage (i.e., below a target cumulative coverage of 10x) and those individuals were sequenced at 1x so that the cumulative coverage on the haplotypes that they carried could be increased (i.e., at or above a target cumulative coverage of 10x). We split the sequencing resources equally between the first and the second stage of AlphaSeqOpt. The number of families of focal individuals targeted in the first stage of AlphaSeqOpt was determined by assuming an average cost of 1400 MU per family, which was equivalent to sequencing all 7 family members (i.e., the focal individual plus its two parents and four grandparents) at 2x. Note that this average cost per family was used only to determine the number of families targeted but that AlphaSeqOpt distributes the total resources differently across different families and different members within these families. AlphaSeqOpt uses phased genotype data when performing its optimisation. In this study we used haplotype libraries built from the true simulated phased marker array genotypes of all 20 chromosomes. We refer to this method as ‘AlphaSeqOpt’.

In addition to these four methods, we also tested one combination of pedigree-and haplotype-based methods and two random methods as controls to separate the contribution of informed methods to imputation accuracy:

##### Combination of pedigree-and haplotype-based methods

We used a combination of the TopSiresAndDams and AlphaSeqOpt methods. To do this, we assumed that sequencing resources were split equally between each method. Thus, 25% of the investment was used for sequencing top sires at 2x, 25% for top dams at 1x, 25% for the focal individuals and their families at variable coverage (stage 1 of AlphaSeqOpt), and the remaining 25% for individuals carrying under-sequenced haplotypes at 1x (stage 2 of AlphaSeqOpt). This method was used as the baseline scenario for comparisons. We refer to this method as ‘Combined’.

##### Random with variable coverage

We randomly selected individuals from the population to be sequenced, while maintaining the same distribution of coverage levels as in the Combined method, i.e., the same number of individuals were sequenced at 1x, 2x, 5x, 15x, or 30x. We refer to this method as ‘RandomVar’.

##### Random at uniform coverage

We used a second random method that selected individuals from the population and sequenced these individuals uniformly at 2x. We refer to this method as ‘RandomUnif’.

#### Test 2: Generation to which the sequenced individuals belong

We tested the effect of the generation to which the sequenced individuals belonged on the imputation accuracy of hybrid peeling. To do this, we created five scenarios per population and level of investment. In each scenario involving populations with real pedigrees, we selected the individuals to sequence randomly but with the constraint that they could only come from a given decile, i.e., individuals were selected from the 0-0.1, 0.2-0.3, 0.4-0.5, 0.6-0.7, or 0.8-0.9 relative positions of the pedigree. The relative position of an individual within a pedigree was defined as its ordinal position after sorting by date of birth. The relative position in the pedigree was used as a proxy for the generation to which an individual belonged in real pedigrees with overlapping generations. In populations with simulated pedigrees, the same procedure was followed but all the sequenced individuals were concentrated into given generations rather than deciles. In this test all selected individuals were sequenced at 2x. We complemented this test by assessing how the distribution of the sequenced individuals selected with the methods tested in Test 1 related to the imputation accuracy across the population.

#### Test 3: Variable or uniform coverage

We compared the use of variable or uniform coverage. To do this, for each level of investment, we took the same individuals that were selected for sequencing at variable coverage with the Combined method in Test 1, and assigned them a uniform coverage. First, we assigned them the average individual coverage generated with the Combined method in Test 1. However, this scenario required a greater total investment than the scenario with variable coverage, because the individuals at higher coverage have proportionally lower cost per x of sequencing. To account for this, we also distributed uniform coverage between the same individuals so that the investment required would equal the investment of the Combined method with variable coverage.

#### Test 4: Number of individuals and sequencing coverage

We tested the effect of the trade-off between the number of sequenced individuals and the coverage at which they were sequenced on the imputation accuracy of hybrid peeling to assess whether it was more beneficial to sequence fewer individuals at a higher coverage or more individuals at a lower coverage. To do this, we created seven scenarios per population and level of investment. In each scenario, sequenced individuals were selected randomly and all the selected individuals were sequenced at either 0.25x, 0.5x, 1x, 2x, 3x, 4x, or 5x. We sequenced as many individuals as the level of investment would allow given the costs of library preparation and sequencing.

### Generation of whole-genome sequence data

After selecting individuals to sequence and sequencing coverage we simulated read counts for each individual and locus from a Poisson-gamma distribution to account for the variability due to sequenceability of each locus and number of reads for each individual at each locus [35]:

1. Sequenceability of each marker locus (*s*_*j*_) was sampled from a Gamma distribution with shape and rate parameter equal to 4, *s*_*j*_ ∼ Gamma(*a* = 4, *b* = 4).
2. For a sequencing coverage of *x*, the number of sequence reads for an individual *i* at a locus *j* (*n*_*i,j*_) was sampled from a Poisson distribution with mean equal to *xs*_*j*_, *n*_*i,j*_ ∼ Poisson(*l* = *xs*_*j*_).
3. The sequence reads were distributed at random between the two alleles of an individual, n_*i,j*,1_ ∼ Binomial(*p* = 0.5, *k* = n_*i,j*_) and n_*i,j*,2_ = n_*i,j*_ – n_*i,j*,1_.

### Hybrid peeling imputation

We performed imputation using hybrid peeling, as implemented in AlphaPeel [1], with the default settings. Hybrid peeling extends the methods of Kerr and Kinghorn [36] for single-locus iterative peeling and of Meuwissen and Goddard [37] for multi-locus iterative peeling to efficiently call, phase and impute whole-genome sequence data in complex multi-generational pedigrees. We performed multi-locus iterative peeling on all available marker array data to estimate the segregation probabilities for each individual. We did not impute the individuals genotyped with LD marker arrays to HD prior to this step. We used the segregation probabilities to perform segregation-aware single-locus iterative peeling for the remaining segregating sites.

To reduce computational demands, we performed single-locus peeling on a random subset of 5,000 non-consecutive SNPs taken from across three chromosomes. While we simulated 20 chromosomes to represent realistic genetic architecture and haplotype diversity (e.g., for the haplotype-based method AlphaSeqOpt), preliminary analyses revealed negligible variation of imputation accuracy across chromosomes and we limited the estimation of imputation accuracy to three chromosomes to reduce the computational requirements of this study.

### Imputation accuracy

We measured imputation accuracy as the individual-wise correlation between true genotypes and imputed dosages. The individual-wise correlation was calculated after correcting for minor allele frequency (MAF), as recommended by Calus et al. [38]. In the context of this study, we found that the relationship between the raw correlation uncorrected for MAF and the dosage corrected for MAF was nearly linear [28]. To facilitate comparison with other studies that report only the uncorrected dosage correlations, we found that the MAF corrected correlations of 0.75, 0.80, 0.85, 0.90, and 0.95 were equivalent to the raw correlations of 0.89, 0.91, 0.93, 0.96, and 0.98 respectively. For comparison of the scenarios in each test, we calculated population-wide imputation accuracy by averaging the individual-wise dosage correlations.

## Results

Imputation accuracy depended on pedigree structure and level of investment in sequencing but it was robust to sequencing strategy as long as there was a sufficient amount of sequence data widely distributed across the generations of a pedigree. In terms of pedigree structure, we found that pedigree depth (number of generations) greatly determined the accuracy of imputation. The gains in imputation accuracy from additional investment in sequencing diminished as population size and level of investment in sequencing increased. The performance of the different methods used for selecting which individuals to sequence depended mostly on to which generation the sequenced individuals belonged, with wider distributions across generations providing more persistent imputation accuracy across the population. There was not a clear benefit to sequencing the selected individuals at variable coverage. Instead, a uniform coverage of around 2x was sufficient to achieve high imputation accuracy.

### Pedigree structure, population size and level of investment

Imputation accuracy greatly increased with pedigree depth (number of generations) but the generation size (number of individuals per generation) had only a small effect on imputation accuracy. Figure 1 shows the imputation accuracy achieved for the populations with different pedigree structures, population sizes, and levels of investment. The results for the populations with simulated pedigrees are plotted against pedigree depth and generation size of each pedigree, while the results for the populations with real pedigrees are plotted against the total number of individuals in the population because they had overlapping generations.

**Figure 1.**
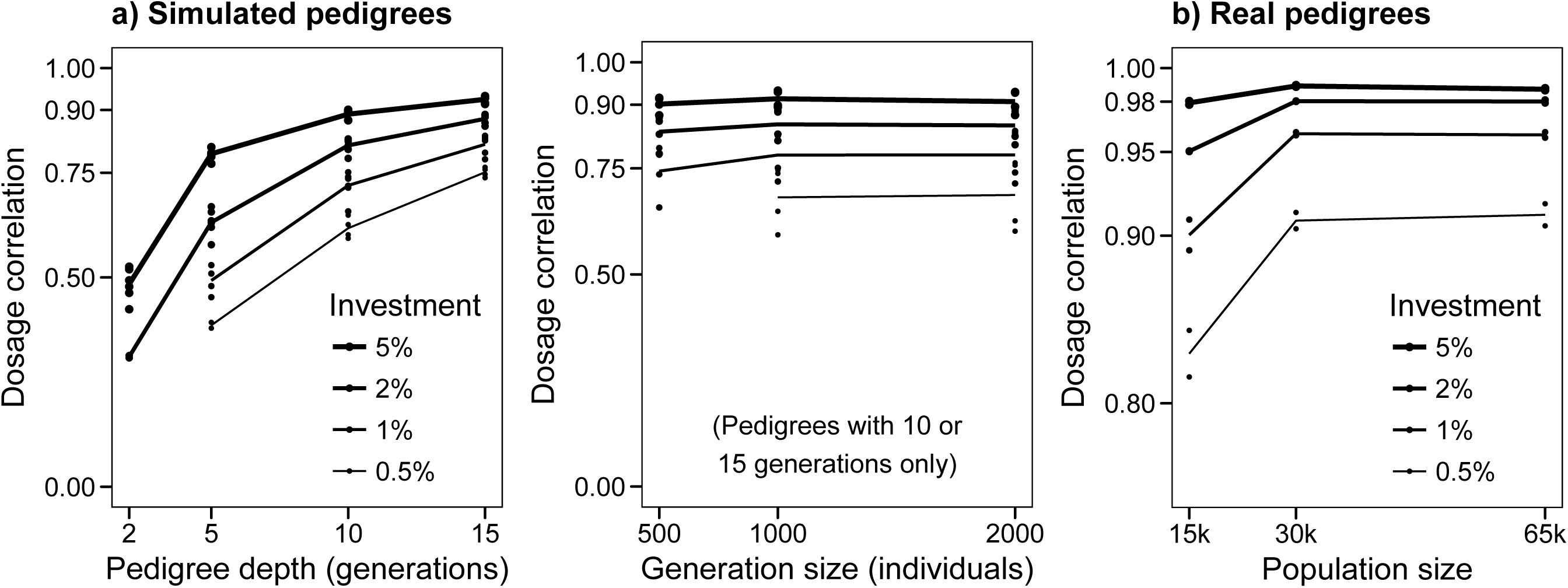
Imputation accuracy for the populations with simulated (a) and real (b) pedigrees by level of investment using the Combined sequencing strategy.

#### Pedigree structure and population size

Populations with very shallow pedigrees had much lower levels of imputation accuracy compared to populations with deep pedigrees, regardless of the generation size. For example, for the maximum level of investment considered (i.e., equivalent to 5% of the population sequenced at 2x), the imputation accuracy averaged across the three populations with simulated pedigrees was 0.48 for 2-generation pedigrees, 0.80 for 5-generation pedigrees, 0.89 for 10-generation pedigrees, and 0.93 for 15-generation pedigrees (Figure 1a). In contrast, quadrupling population size by increasing the generation size from 500 to 2,000 individuals per generation only increased imputation accuracy from 0.44 to 0.52 in the 2-generation pedigrees, from to 0.81 in the 5-generation pedigrees, and from 0.92 to 0.93 in the 15-generation pedigrees, while it did not change in the 10-generation pedigrees.

The size of the populations with simulated pedigrees ranged from 1,000 to 30,000 individuals. In the populations with simulated pedigrees, population size affected the accuracy of imputation accuracy to the extent that deeper pedigrees were larger. The populations with real pedigrees were larger than most of the simulated pedigrees, with approximately 15,000 to 65,000 individuals, and encompassed approximately 20 overlapping generations. The imputation accuracy achieved for the populations with real pedigrees was higher than those of populations with simulated pedigrees of a similar size (Figure 1b).

#### Level of investment

The level of investment in sequencing also had a strong impact on imputation accuracy, particularly for the smallest populations and for the lowest levels of investment. There were diminishing returns for large populations and large levels of investment, probably because high imputation accuracy was already achieved in these scenarios and differences became less noticeable. To achieve the same imputation accuracy, populations with shallow pedigrees required a proportionally greater sequencing effort than populations with deep pedigrees. For example, to achieve dosage correlations of ∼0.80, 5% of the individuals from the 5-generation populations needed to be sequenced but only 2% from the 10-generation populations or 1% from the 15-generations populations.

#### Imputation accuracy across generations

Imputation accuracy differed across generations of the pedigree. Imputation accuracy was low in the first generations and then increased in subsequent generations until it stabilised. Figure 2 shows the average imputation accuracy achieved for the individuals in each generation of the simulated pedigrees. With the maximum level of investment in sequencing (Figure 2a,b,d), imputation accuracy seemed to plateau at generation 4. At lower levels of investment in sequencing, imputation accuracy increased more slowly and did not plateau until a later generation. These patterns were independent of generation size, but imputation accuracy by generation was always greater in populations with deep pedigrees than with shallow pedigrees. A similar pattern was observed in the real pedigrees (results not shown), where we observed reduced imputation accuracy for the individuals in the first generations.

**Figure 2.**
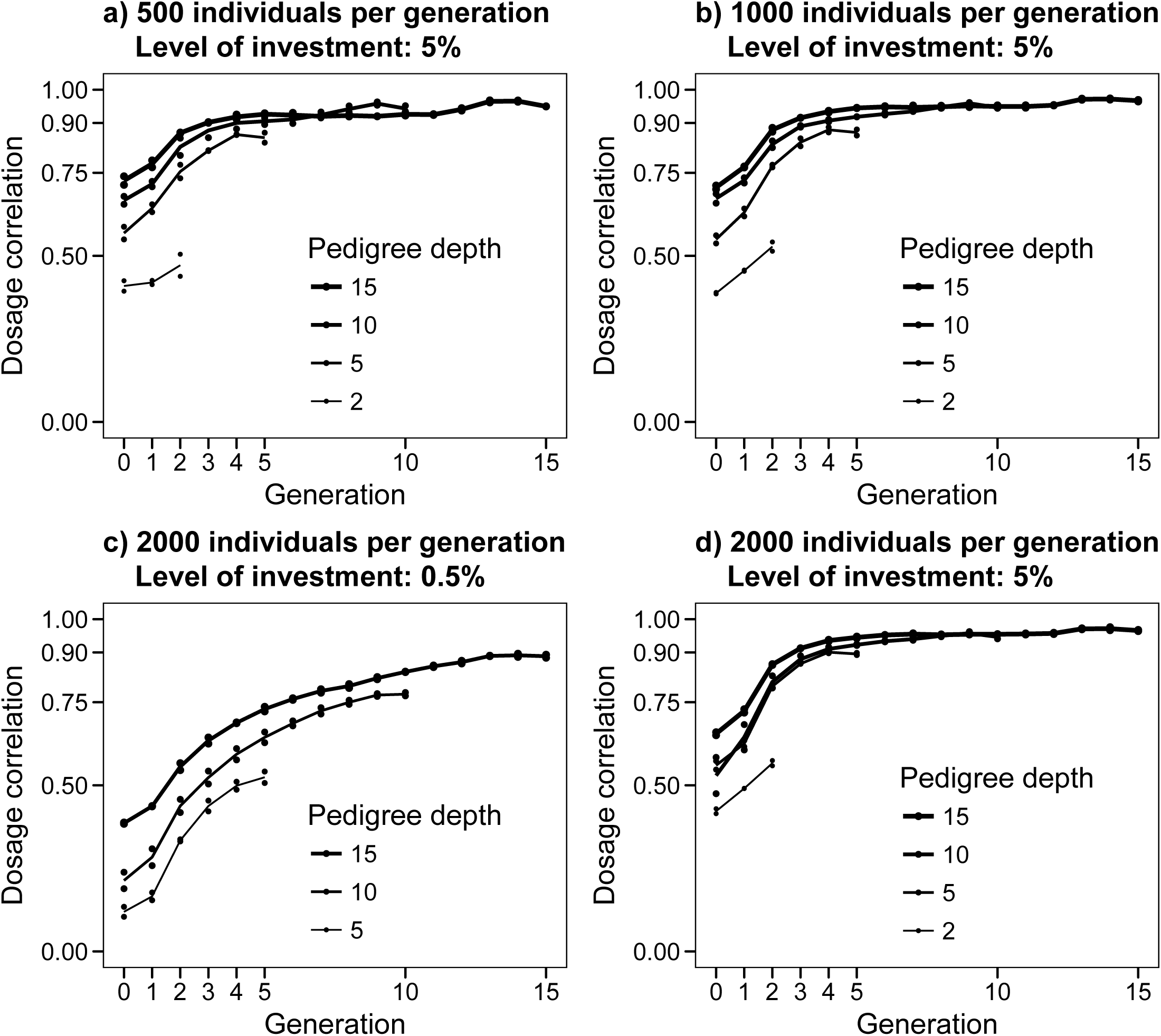
Imputation accuracy by generation for different simulated pedigrees and levels of investment. The sequenced individuals and their coverage were selected using the Combined method.

The lower imputation accuracy in the first generations can be explained by insufficient information from immediate ancestors to accurately estimate the segregation probabilities on which hybrid peeling relies [1,28] (i.e., their parents and grandparents were unknown or did not have any marker array data). In light of these results, the populations with simulated pedigrees with 2 or 5 generations were not considered further. Imputation accuracy of the populations with simulated pedigrees in the remaining tests was assessed using only individuals from generation 4 onwards, and imputation accuracy of populations with real pedigrees was assessed after discarding the first 20% individuals of the pedigree.

#### Imputation accuracy in populations with overlapping generations

Imputation accuracy of the populations with real pedigrees after discarding the first 20% individuals (Figure 1b) ranged from 0.98 to 0.99 with the maximum level of investment considered (i.e., equivalent to 5% of the population sequenced at 2x), and displayed no observable trend for population size. The imputation accuracy with the minimum level of investment (i.e., equivalent to 0.5% of the population sequenced at 2x) was already high for the populations with 30k and 65k individuals (0.91) but lower for the population with 15k individuals (0.83). As level of investment increased, the difference between the 15k population and the two larger ones decreased from 0.08 (with a level of investment of 0.5%) to 0.06 (with level of investment of 1%), 0.03 (with a level of investment of 2%) and only 0.01 (with a level of investment of 5%). There were no noticeable differences in imputation accuracy between the populations with 30k and 65k individuals, and both achieved imputation accuracy over 0.98 with levels of investment of 2% or more.

### Test 1: Method for selecting sequenced individuals and coverage

Imputation accuracy was consistently high for many of the methods tested, although the ranking of best performing methods depended on the level of investment. In general, the pedigree-and haplotype-based methods Combined, TopSiresAndDams, AlphaSeqOpt, and PedConnect were amongst the best performing methods, together with the methods based on random selection of individuals (RandomVar and RandomUnif). Figure 3 shows the imputation accuracy achieved with each method. To simplify, we only show results from the populations with real pedigrees and the simulated pedigrees with 10 or 15 generations and 2,000 individuals per generation and for the two most extreme levels of investment (i.e., equivalent to 0.5% or 5% of the population sequenced at 2x).

**Figure 3.**
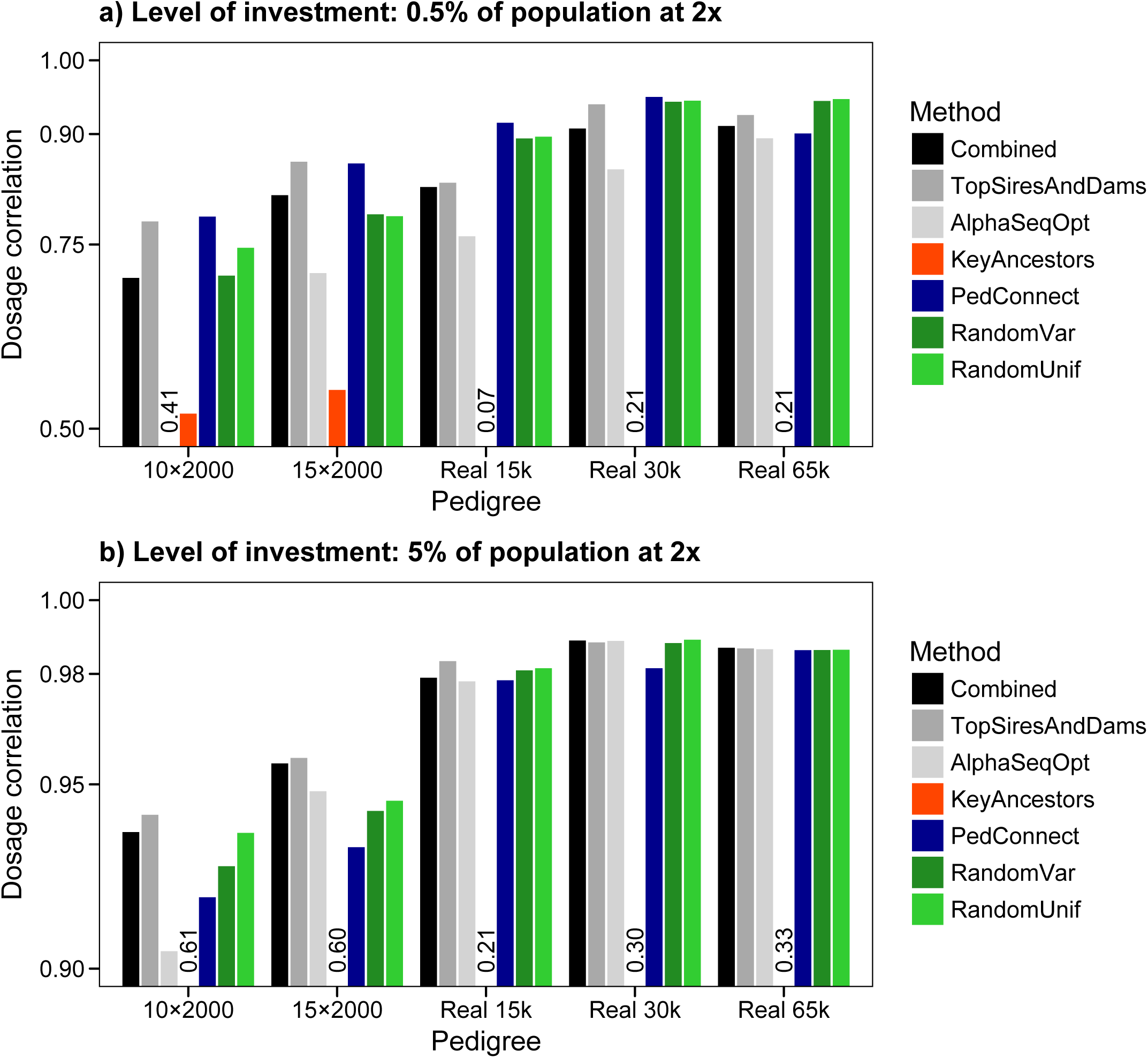
Imputation accuracy obtained using seven methods for selecting individuals to sequence and sequencing coverage, with an investment of either 0.5% or 5% of the population sequenced at 2x. Bars below axis limit are represented with a number.

At high levels of investment the methods Combined, TopSiresAndDams, AlphaSeqOpt, PedConnect, RandomVar, and RandomUnif gave similar high imputation accuracy, of 0.97 to 0.99, for the populations with real pedigrees. At low levels of investment either PedConnect or the random methods RandomVar and RandomUnif generally performed better for the populations with real pedigrees than Combined, TopSiresAndDams, and AlphaSeqOpt, with differences being more noticeable in small populations. For the populations with simulated pedigrees, TopSiresAndDams was the best performing method, although there was more variability in the ranking of the methods than for the populations with real pedigrees. In the populations with simulated pedigrees, the PedConnect method performed similarly to TopSiresAndDams when the level of investment was low but its performance dropped when the level of investment was high. In some of the populations with simulated pedigrees, AlphaSeqOpt produced much lower imputation accuracy than the best performing methods, although this was not observed in the populations with real pedigrees. The KeyAncestors method performed consistently worse than all other methods.

### Test 2: Generation to which sequenced individuals belong

Imputation accuracy in each generation of the pedigree depended on the generation to which the sequenced individuals belonged. Figure 4 shows the position of the sequenced individuals within the pedigree, their sequencing coverage, and the imputation accuracy obtained for the individuals along the pedigree for the methods included in Test 1. For simplicity, we only plotted results of the population with a real pedigree with 30k individuals and a level of investment equivalent to 2% of the population sequenced at 2x.

**Figure 4.**
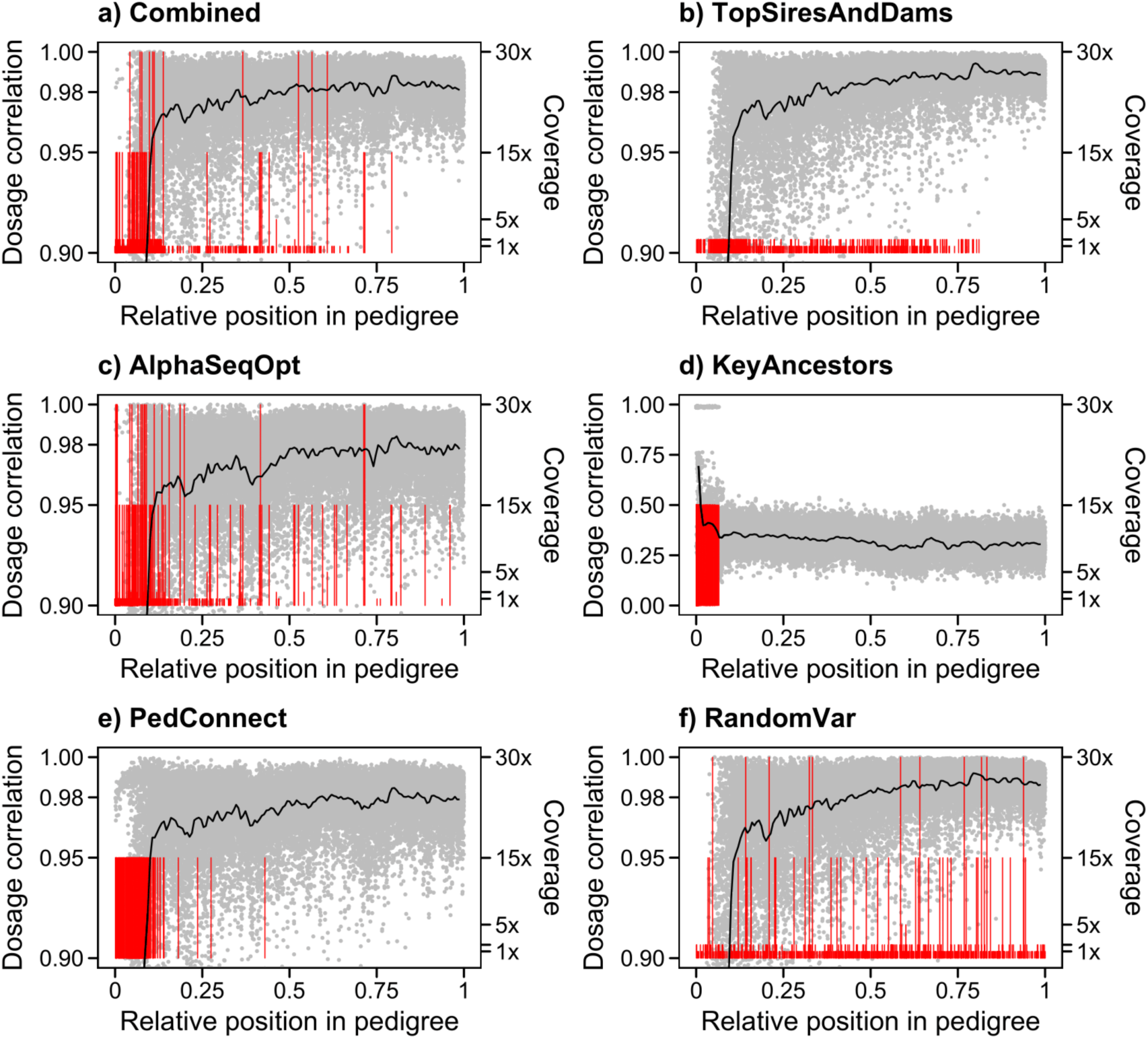
Imputation accuracy obtained along the pedigree with different methods for selecting individuals to sequence and sequencing coverage. Results are for a population with a real pedigree with 30k individuals with investment equivalent to 2% of the population sequenced at 2x. The vertical red lines represent the position of the sequenced individuals and their length represents sequencing coverage (1, 2, 5, 15, or 30x). The grey dots represent the imputation accuracy obtained for each individual. The black line is the moving average. The profile obtained for RandomUnif was very similar to that with the RandomVar method, but with more individuals all at 2x.

The KeyAncestors method targeted only individuals from the first generations and imputation accuracy across all generations was low. The PedConnect method also targeted individuals from the first generations but the sequenced individuals spanned a wider portion of the generations in the pedigree. Thus, for PedConnect imputation accuracy rapidly increased from very low for the first generations to over 0.95 for the generations that were immediately posterior to the bulk of sequenced individuals and then it continued to increase slowly in subsequent generations. For the rest of methods we found the same pattern of the imputation accuracy along the generations of the pedigree, with slight differences in imputation accuracy in early or late generations depending on the position of the sequenced individuals. The individuals selected for sequencing by the Combined and TopSiresAndDams methods were more widely distributed across the generations but with a greater concentration in early generations. The TopSiresAndDams method had less dispersion of individual-wise imputation accuracy than other methods such as AlphaSeqOpt and PedConnect. The distribution of the sequenced individuals across the generations selected with the AlphaSeqOpt method did not differ much from that of the Combined method but it had a greater number of individuals sequenced at high coverage, most of which were from early generations. Imputation accuracy with the AlphaSeqOpt method was lower than with the Combined method across all generations of the pedigree. With the RandomVar and RandomUnif (not shown) methods the sequenced individuals were more evenly distributed across the generations of the pedigree and imputation accuracy followed a similar pattern to the Combined and TopSiresAndDams methods but with lower imputation accuracy in the first generations.

The generation of the pedigree to which the sequenced individuals belonged greatly determined the imputation accuracy in the rest of generations. That effect was more easily characterized when all the sequenced individuals were concentrated in a single decile or generation of the pedigree. Figure 5 shows the imputation accuracy obtained along the pedigree when the sequenced individuals belonged to a single decile of the real pedigrees. Figure 6 shows the imputation accuracy obtained along the pedigree when the sequenced individuals belonged to a single generation of the simulated pedigrees.

**Figure 5.**
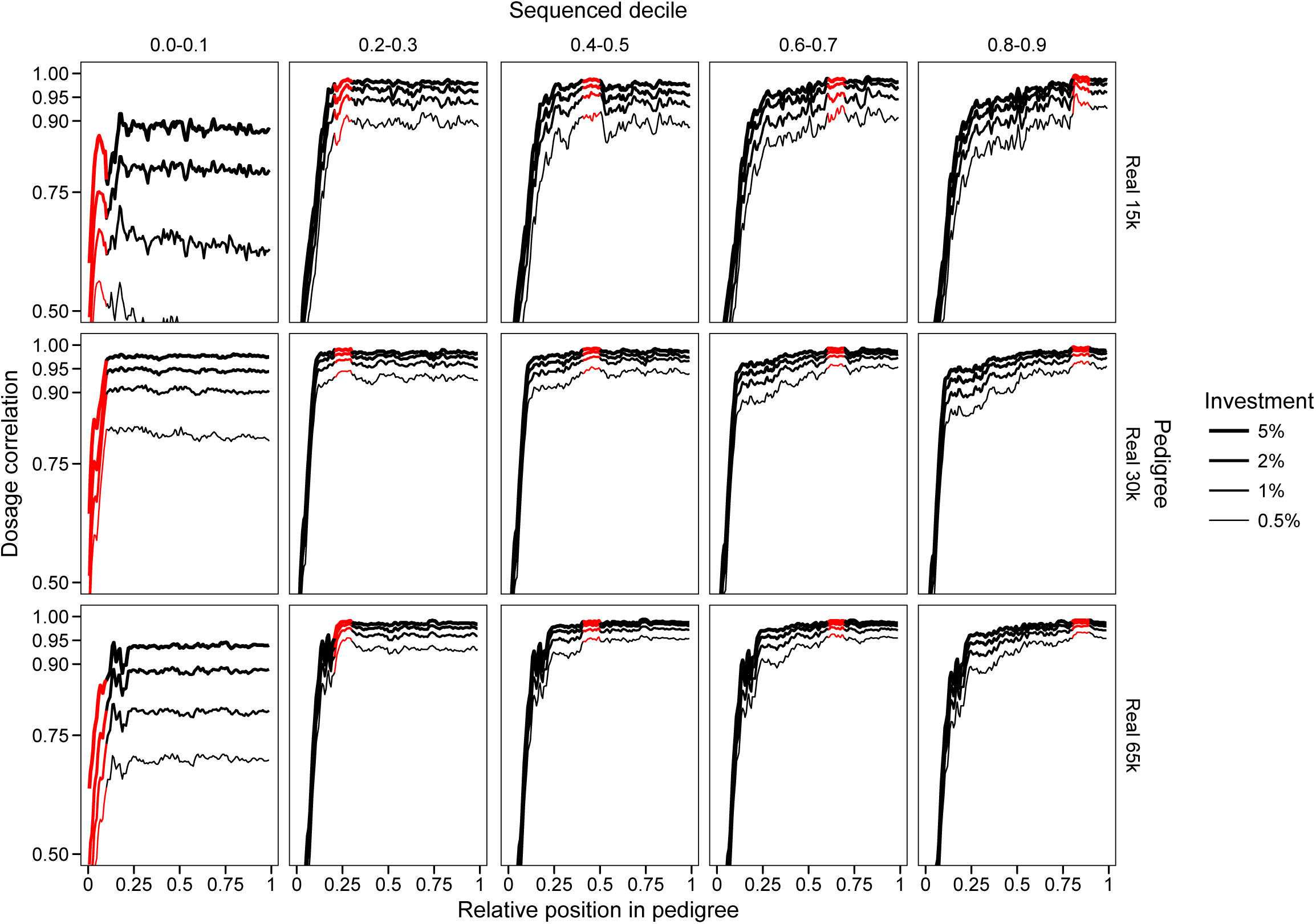
Imputation accuracy obtained along the real pedigrees when the sequenced individuals are concentrated in a given decile of the pedigree. The red segment represents the decile where the sequenced individuals were located. Within that decile, sequenced individuals were chosen randomly and sequenced at 2x. Different levels of investment were considered.

**Figure 6.**
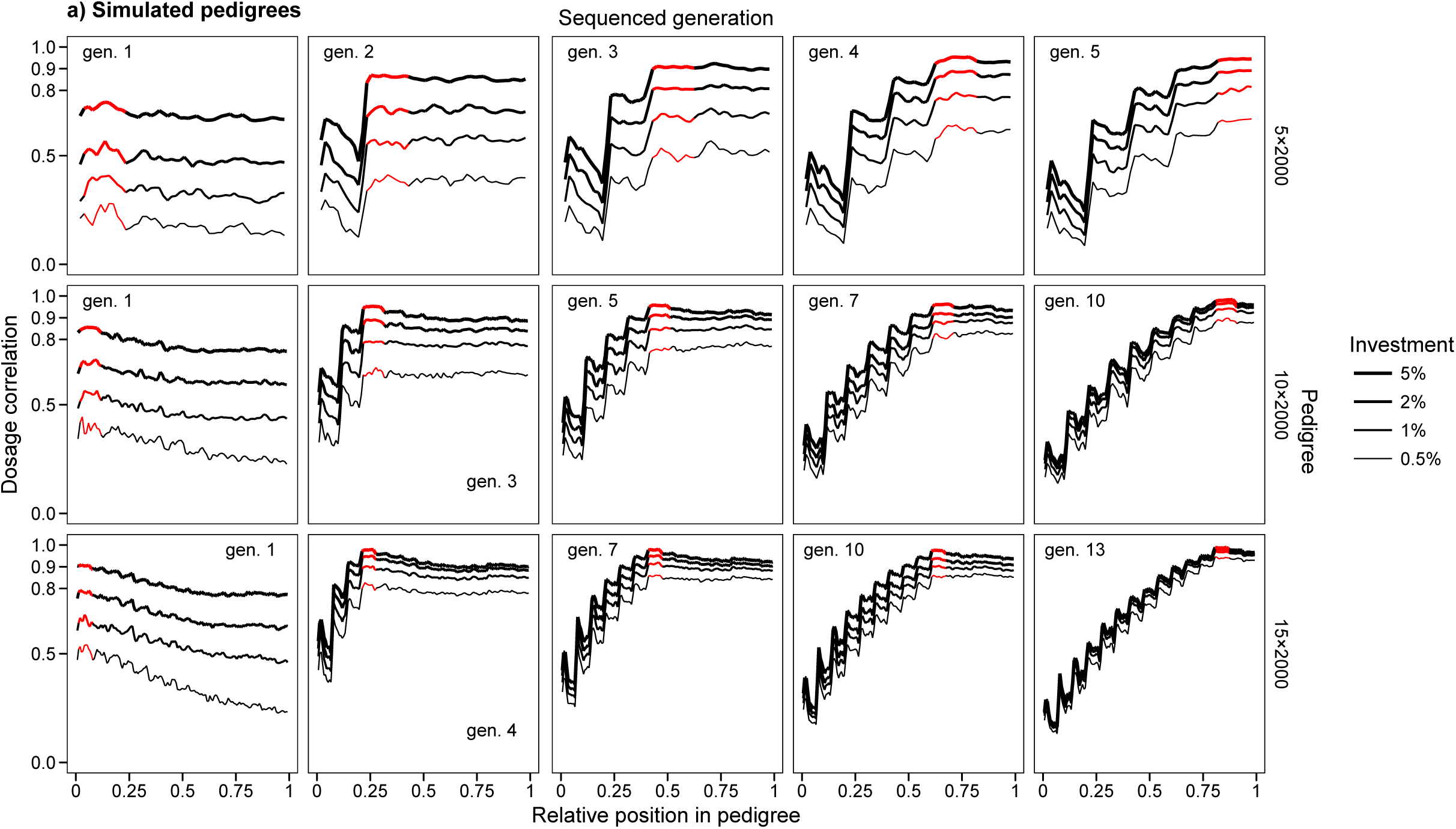
Imputation accuracy obtained along the simulated pedigrees when the sequenced individuals are concentrated in a given generation. The red segment represents the generation where the sequenced individuals were located. Within that generation, the sequenced individuals were chosen randomly and sequenced at 2x. Different levels of investment were considered. Base generation was counted as generation 0.

Sequencing the individuals from the first decile resulted in very poor imputation accuracy across the population. However, sequencing in any of the deciles from the third decile onwards provided high imputation accuracy in that decile and the imputation accuracy largely persisted in the subsequent deciles. Theoretically, some decay of imputation accuracy should be expected as the imputed individuals from the subsequent deciles became more distant from the sequenced individuals but in our test this decay was mostly negligible in the populations with real pedigrees. However, decay was observed in the population with simulated pedigrees (Figure 6). In contrast, imputation accuracy in deciles previous to the one to which the sequenced individuals belonged had a more pronounced decay as the imputed individuals became more distant from the sequenced individuals, until imputation accuracy decayed drastically for the first deciles in the pedigree. We observed the same pattern in the populations with simulated pedigrees.

### Test 3: Variable or uniform sequencing coverage

There was no clear benefit to using variable, as opposed to uniform, levels of sequencing coverage. Figure 7 shows the imputation accuracy achieved when the same individuals selected with the Combined method were sequenced at variable coverage or at a uniform coverage. Table 2 compares the number of individuals sequenced, sequencing coverage, and investment required with the Combined method with variable coverage or with uniform coverage across the same sequenced individuals.

**Table 2.** Number of sequenced individuals, distribution of sequencing coverages, total coverage, and total investment for the different populations with the Combined method with variable or uniform coverage.

| Pedigree | Variable coverage<sup>a</sup> |  |  |  |  |  |  |  | Uniform coverage at equal total coverage<sup>b</sup> |  | Uniform coverage at equal investment<sup>c</sup> | Coverage of 2x at equal investment<sup>d</sup> |
| --- | --- | --- | --- | --- | --- | --- | --- | --- | --- | --- | --- | --- |
|  | Number of individuals sequenced |  |  |  |  |  | Total coverage (x) | Investment (MU<sup>e</sup>) | Individual coverage (x) | Investment (MU<sup>e</sup>) | Individual coverage (x) | Number of individuals sequenced |
|  | Total | 1x | 2x | 5x | 15x | 30x |  |  |  |  |  |  |
| 2×2000 | 223 | 161 | 47 | 0 | 8 | 7 | 585 | 38,670 | 2.62 | 55,720 | 1.67 | 193 |
| 5×2000 | 561 | 403 | 119 | 1 | 27 | 11 | 1381 | 95,450 | 2.46 | 132,920 | 1.63 | 477 |
| 10×2000 | 1130 | 813 | 237 | 4 | 56 | 20 | 2747 | 191,720 | 2.43 | 264,960 | 1.62 | 958 |
| 15×2000 | 1703 | 1223 | 355 | 11 | 89 | 25 | 4073 | 288,350 | 2.39 | 393,960 | 1.62 | 1441 |
| Real 15k | 877 | 632 | 182 | 3 | 50 | 10 | 2061 | 147,060 | 2.35 | 199,960 | 1.60 | 735 |
| Real 30k | 1642 | 1245 | 281 | 9 | 76 | 31 | 3922 | 273,910 | 2.39 | 379,440 | 1.59 | 1369 |
| Real 65k | 3742 | 2711 | 773 | 37 | 156 | 65 | 8732 | 629,450 | 2.33 | 848,240 | 1.60 | 3147 |

<sup>a</sup> Results obtained with the Combined method with an investment equivalent to 5% of the population sequenced at 2x.
<sup>b</sup> The total coverage obtained with the Combined method distributed uniformly across the same individuals.
<sup>c</sup> The investment allocated with the Combined method distributed uniformly across the same individuals.
<sup>d</sup> The investment allocated with the Combined method distributed uniformly across as many individuals as can be sequenced at 2x.
<sup>e</sup> Monetary units.

**Figure 7.**
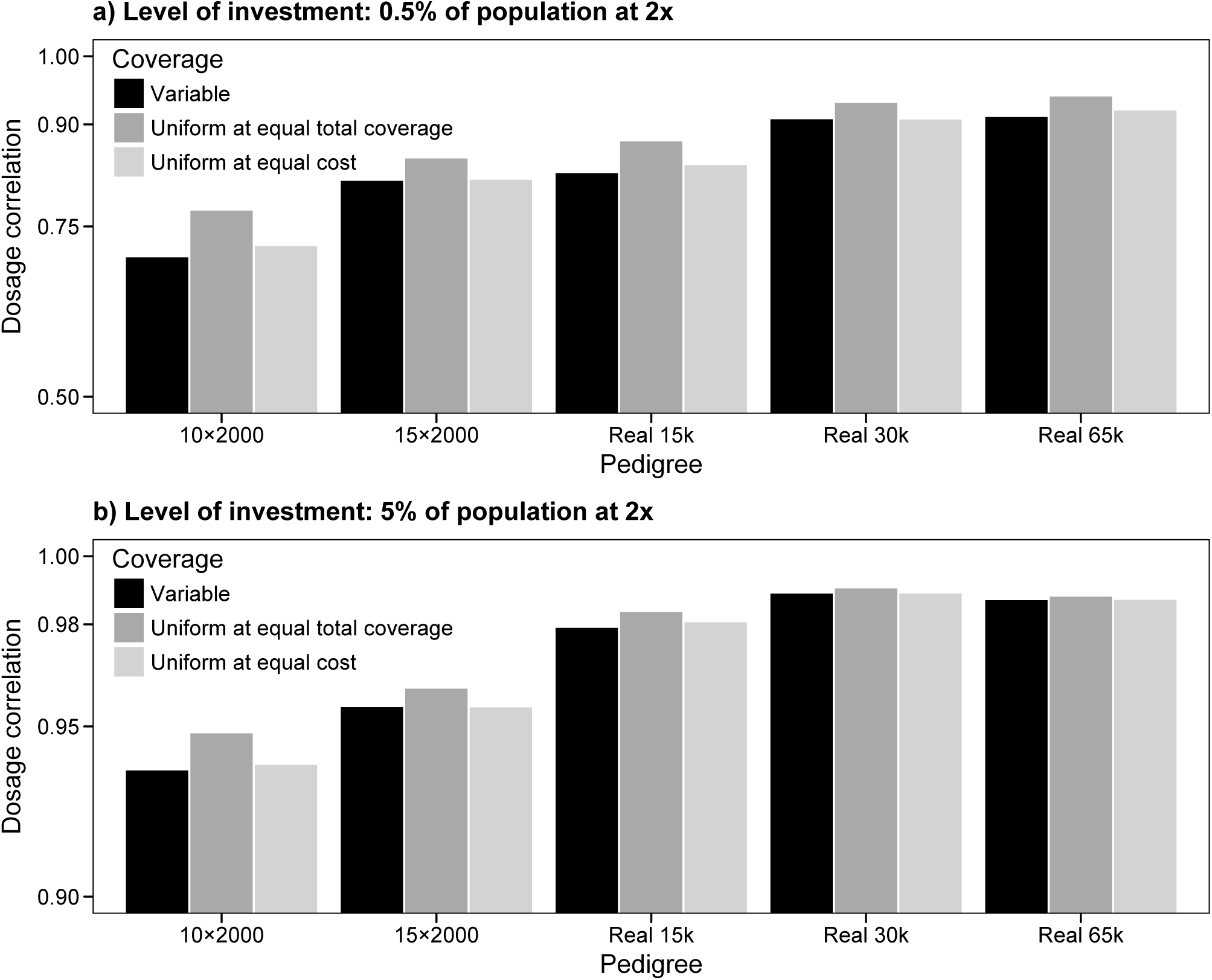
Imputation accuracy with different coverage distribution across the sequenced individuals, with an investment of either 0.5% or 5% of the population sequenced at 2x. The sequenced individuals were selected using the Combined method. The sequencing coverage was either variable or uniform, in which case either the total coverage or the total cost was distributed uniformly across the sequenced individuals.

For the same amount of total coverage it was more beneficial to distribute the sequencing resources uniformly across the same set of sequenced individuals. At the same amount of total coverage, imputation accuracy when sequencing at uniform coverage was generally greater than with variable coverage, by between 0.02 and 0.07 when the level of investment was 0.5% but by only up to 0.01 when the level of investment was 5%. However, sequencing the same amount of total coverage using uniform coverage across the same set of sequenced individuals would incur in a 40% greater sequencing cost due to the assumed non-linear cost structure. Therefore, we considered an additional scenario of uniform coverage where investment equalled the investment of the scenario of variable coverage. At equal investment, the realized total coverage decreased by 35% when the individuals were sequenced at uniform coverage compared to the scenario of variable coverage. Imputation accuracy in the scenario of variable coverage at equal investment increased by up to 0.02 compared to the scenario of variable coverage when the level of investment was 0.5%, although it did not change noticeably when the level of investment was 5%. At equal investment, if all the selected individuals were sequenced at 2x, on average 15% fewer individuals could be sequenced compared to the scenario with variable coverage, due mostly to the cost associated with library preparation.

### Test 4: Number of individuals and sequencing coverage

When all the individuals were sequenced at a uniform coverage, the sequencing coverage that produced the highest imputation accuracy was around 2x, especially at low levels of investment. Figure 8 shows the imputation accuracy achieved for the populations with real pedigrees with different levels of investment when all the individuals were sequenced at different levels of coverage. In all three populations there was a small reduction in imputation accuracy when coverage was 1x or less. That reduction was on average 0.03 (max: 0.08) with a level of investment of 0.5% and only up to 0.01 with a level of investment of 5%. Similarly, there was a less pronounced reduction of imputation accuracy when sequencing coverage increased. This reduction was on average 0.01 (max: 0.03) when the level of investment was 0.5%, but unnoticeable at the highest levels of investment.

**Figure 8.**
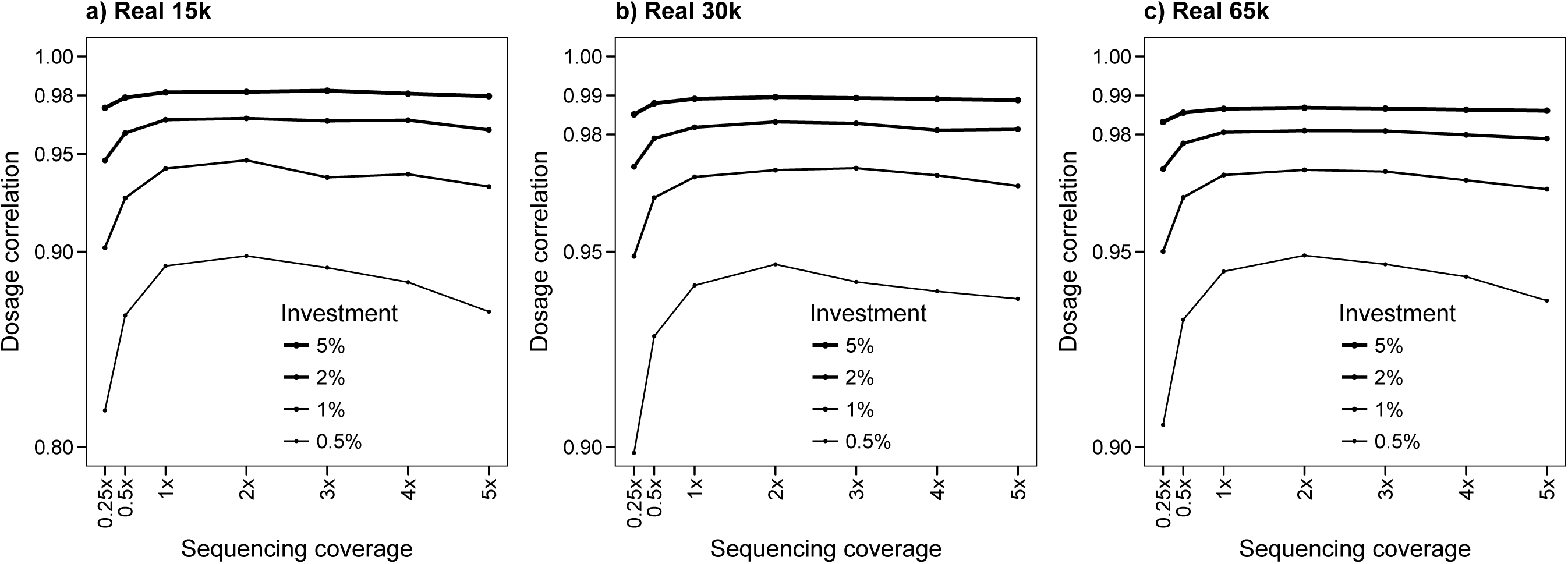
Imputation accuracy in real pedigrees when as many individuals as affordable were sequenced at different levels of coverage and levels of investment. The sequenced individuals were selected randomly.

## Discussion

The choice of sequencing strategy and method for selecting individuals to sequence should be tailored to the properties of the methods that will subsequently be used for variant discovery and for imputation. We performed a series of simulations to assess how the sequencing strategy can affect the imputation accuracy of hybrid peeling for a set of populations with simulated and real pedigrees with different pedigree structures and population sizes. The results show that: (i) imputation accuracy depends largely on pedigree depth; (ii) hybrid peeling is robust to the method used for selecting which individuals to sequence and at which coverage, as long as the number of individuals sequenced is large and the sequenced individuals are distributed widely across all generations in the pedigree; and (iii) a sequencing coverage of 2x for all the sequenced individuals is sufficient to achieve high imputation accuracy when the number of sequenced individuals is large.

In what follows we will discuss: (i) the impact of pedigree structure; (ii) the impact of population size and level of investment; (iii) the impact of the selection of individuals to sequence; (iv) the impact of the sequencing coverage; (v) the recommendations for sequencing strategies suited for pedigreed populations when using whole-population imputation with hybrid peeling; and (vi) the limitations of the study.

### Impact of pedigree structure

Pedigree structure determined overall imputation accuracy. Pedigree depth was the factor with the greatest effect on imputation accuracy, while generation size did not have a noticeable effect. This occurred because during peeling information is transmitted ‘vertically’ across generations (from parents to progeny and vice versa) and transmitted ‘horizontally’ to contemporary individuals only if they are connected via a relative in another generation (e.g., a common ancestor). In practice, real livestock pedigrees have overlapping, rather than discrete, generations with many closely related individuals, which typically leads to high imputation accuracy.

### Impact of population size and level of investment

Gains in imputation accuracy from additional investment in sequencing resources diminished as population size and levels of investment increased. This occurred because the high connectedness between individuals in livestock populations favours the diffusion of information from the sequenced individuals to distant relatives, to the point that information from additional sequenced individuals was increasingly redundant. For livestock populations with a size of similar order of magnitude as the real pedigrees that we tested our results suggest that the minimum amount of sequence data required for high imputation accuracy should be the equivalent of sequencing approximately 2% of the population at 2x. However, smaller populations require proportionally greater investment to achieve the same imputation accuracy as larger populations.

### Impact of the selection of individuals to sequence

#### Impact of the generation to which the sequenced individuals belong

One of the main factors that determined imputation accuracy across the population was the generation of the pedigree to which the sequenced individuals belonged. We observed two distinct trends:

##### Sequencing allocated to individuals from the first two generations or that do not have genotype data for at least a couple of generations of ancestors

Sequencing such individuals, even at high coverage, failed to produce good imputation accuracy for the rest of the population. This may seem counterintuitive at first but is explained by the fact that the accuracy of the segregation probabilities for these individuals, which are used in hybrid peeling, was low due to insufficient information from their immediate ancestors (i.e., their parents and grandparents were unknown or did not have any marker array data) [1,28]. The poor accuracy of their segregation probabilities hindered the transmission of information between the first generation and the subsequent ones.

##### Sequencing allocated to individuals that have genotype data for at least a couple of generations of ancestors

Sequencing such individuals produced high imputation accuracy for the rest of the population. However, in these situations there was a decay in imputation accuracy as the pedigree distance between the sequenced individuals and those to be imputed increased. This decay was more pronounced when individuals in earlier generations were imputed from sequenced individuals in later generations than vice versa, because descendants are generally less informative for imputation than ancestors. It was of note that in the populations with real pedigrees we did not observe a decay when individuals in later generations were imputed from sequenced individuals in earlier generations, probably because the high connectedness between the individuals in the real pedigrees enhanced the persistence of the imputation accuracy.

#### Performance of the methods for selecting individuals to sequence

As a consequence of the impact of the generation to which the sequenced individuals belong, the distribution of the sequenced individuals across the pedigree determined the performance of the methods for selecting which individuals to sequence.

##### The TopSiresAndDams method

Targeting the sires and dams that contributed more progeny and grandprogeny to the population was the most effective strategy at the levels of investment that we recommend (i.e., equivalent to sequencing at least around 2% of the population at 2x). The information from these sires and dams was directly transmitted to a large number of descendants widely distributed across the generations of a pedigree. The TopSiresAndDams method also favoured slightly more balanced imputation accuracy along the generations of the pedigree, because sires and dams in early generations are otherwise more difficult to impute from their descendants.

##### The PedConnect method

The PedConnect method was more effective than the TopSiresAndDams method with low levels of investment but not with high levels of investment. The rationale for using the PedConnect method was to maximise the degree of connectedness between the sequenced and imputed individuals to maximize the diffusion of information. Sequencing the individuals selected with this method at high coverage was effective when the level of investment was insufficient for sequencing enough individuals at low coverage for hybrid peeling to probabilistically combine their data. Nonetheless, with a sufficient level of investment, sequencing more individuals at low coverage with the TopSiresAndDams method provided higher imputation accuracy with less dispersion of individual-wise imputation accuracy than the PedConnect method.

##### Other methods

The performance of other methods, such as AlphaSeqOpt or the haplotype-based methods proposed by Druet et al. [7] or Bickhart et al. [16], will largely depend on how they distribute sequencing across the generations of a pedigree.

### Impact of the sequencing coverage

Uniform low coverage at around 2x provided the highest imputation accuracy. No obvious benefit was observed for sequencing individuals at variable coverage. The rationale for assigning variable coverage to the sequenced individuals within a population was to ensure that some families had enough resolution to be able to accurately phase the common haplotypes that they carry [17], so that the phased haplotypes could later be used to impute individuals that shared the same haplotypes [18]. Our results showed that hybrid peeling is able to probabilistically combine low-coverage genotype data across distant relatives, and therefore explicitly calling and phasing specific individuals in the population is not needed.

Additionally, sequencing everybody at a low uniform coverage allows the sequencing of a larger number of individuals. Increasing the number of sequenced individuals results in higher imputation accuracy because this strategy generates data in more individuals from which information can propagate to their relatives. This observation was consistent with empirical observations on real data which showed that the number of individuals with sequence data, rather than the cumulative sequencing coverage, determined imputation accuracy at each variant site [28]. However, when coverage is too low, imputation accuracy falls because many sites are not sufficiently covered with sequence reads to be informative for heterozygote genotypes. Thus the optimal coverage that produced the greatest imputation accuracy was 2x. These results are consistent with results from previous studies. VanRaden et al. [25] showed that, for the same total amount of coverage, reference panels with individuals sequenced at a coverage of 2x to 4x provided the greatest imputation accuracy when using findhap4.

### Recommendations for sequencing strategies for pedigreed populations

Based on our results, we recommend the following sequencing strategy for pedigreed livestock populations when using hybrid peeling for imputation:

i. Maximise the number of sequenced individuals while avoiding sequencing individuals below 2x. There was no clear benefit of sequencing individuals at a higher coverage if the number of sequenced individuals was large. Coverage below 2x is not informative enough for heterozygote loci.
ii. Target for sequencing the sires and dams that contributed more progeny and grandprogeny to the population. This is a simple but effective strategy in livestock pedigrees, where most individuals are direct descendants of a relatively low number of sires and dams. This method selects individuals that are widely distributed across all generations of the pedigree. This distribution of sequenced individuals improves the persistence of imputation accuracy both for early and late generations.
iii. Use deep pedigrees and favour sequencing of individuals that have parents and grandparents with marker array information. The parents and grandparents of the sequenced individuals do not need to be sequenced themselves, but they should be genotyped with marker arrays to enable accurate estimation of segregation probabilities of the sequenced individuals.
iv. Sequence at least around 2% of the individuals in the population if using a coverage of 2x. Smaller populations require proportionally greater investment to achieve the same imputation accuracy as larger populations. Our results can be used as guidance for decision-making on level of investment in pedigrees of similar size with very closely related individuals.

### Limitations of the study

#### Use of simulated data instead of real data

Excessive costs make it impossible to generate sequence data to empirically evaluate several alternative sequencing strategies. Therefore we used simulations. In general, our simulated results were in line with observations in real data [28].

#### Variant discovery

Besides imputation, variant discovery is the other main process affected by the sequencing strategy. We have not tested the impact of the sequencing strategy on variant discovery. However, as with imputation, sequencing at low coverage is a favourable sequencing strategy for variant discovery, especially for increasing discovery rate of rare variants [39,40]. We previously estimated that approximately 75% of the variants discovered in 26 individuals sequenced at 30x can be discovered with the same individuals sequenced at 2x [41]. This discovery rate is expected to further increase with the large sample sizes that are needed for imputation. Other studies have placed the optimum sequencing coverage for variant discovery at a higher coverage of 10 to 12x [7]. Sequencing a subset of individuals at high coverage may improve the variant discovery rate as well as provide a validation set for variants discovered with low-coverage sequence data [28]. For variant discovery, haplotype-based methods such as the second stage of AlphaSeqOpt [18], which is designed to assign low coverage to individuals that carry haplotypes that are not well represented enough in the sequenced individuals, may help in reducing redundancy between the sequenced individuals and increase the number of variants discovered.

#### Classification of variants by minor allele frequency

We found that a sequencing coverage of 2x provided the greatest imputation accuracy. Druet et al. [7] previously estimated that the best imputation accuracy with Beagle [22] was obtained at a higher coverage of 8x for all minor allele frequency categories except for rare variants, for which 2x was the optimal coverage. In our study we did not assess imputation accuracy based on minor allele frequency of the variants.

#### Impact of the marker array genotyping strategy

The strategy for genotyping the individuals using marker arrays (i.e., how many individuals and at which density) is likely also very influential on the imputation accuracy of hybrid peeling. We did not test any of these strategies. There exists a body of work on this topic [20,42–44] and current genotyping practices such as those of breeding programs with genomic selection can be followed.

## Conclusion

Suitable sequencing strategies for subsequent imputation with hybrid peeling involve targeting at least around 2% of the population for sequencing at a uniform coverage around 2x. Hybrid peeling was robust to the method used for selecting which individuals to sequence, as long as the number of individuals sequenced was large and the sequenced individuals were distributed widely across all generations in the pedigree, preferably from the third generation of the pedigree onwards, to improve persistence of imputation accuracy in both early and late generations. Sequencing the sires and dams that contributed more progeny and grandprogeny in the population was a simple but effective strategy. The sequencing strategy that we recommend would be beneficial for generating whole-genome sequence data in populations with deep pedigrees of closely related individuals.

## Ethics approval and consent to participate

Not applicable.

## Consent for publication

Not applicable.

## Availability of data and material

The simulated pedigrees used in this study are available upon request. The real pedigrees used in this study are derived from the PIC breeding programme and not publicly available.

## Authors’ contributions

RRF designed the study; RRF and AW performed the analyses; RRF wrote the first draft with input from JMH; AW, GG and AJM assisted in the interpretation of the results and provided comments on the manuscript. All authors read and approved the final manuscript.

## Acknowledgements

This work has made use of the resources provided by the Edinburgh Compute and Data Facility (ECDF) (http://www.ecdf.ed.ac.uk/).

